# Disrupted Lipid Homeostasis as a Pathogenic Mechanism in *ABCA7*-Associated Alzheimer’s Disease Risk

**DOI:** 10.1101/2025.09.03.673792

**Authors:** Younji Nam, Brooke A. DeRosa, Aura M. Ramirez, Biniyam A. Ayele, Patrice Whitehead-Gay, Larry D. Adams, Charles G. Golightly, Takiyah D. Starks, Mayra Juliana Laverde-Paz, Holly N. Cukier, Rufus Akinyemi, Fred Sarfo, Albert Akpalu, Michael L. Cuccaro, Scott M. Williams, Allison Caban-Holt, Christiane Reitz, Jonathan L. Haines, Goldie S. Byrd, Farid Rajabli, Derek M. Dykxhoorn, Juan I. Young, Jeffery M. Vance, Margaret A. Pericak-Vance

**Affiliations:** John P. Hussman Institute for Human Genomics, University of Miami Miller School of Medicine, 1501 NW 10th Avenue, Miami, FL 33136, USA; Department of Epidemiology and Medical Statistics, College of Medicine, University of Ibadan, Ibadan 200212, Nigeria; Department of Social Sciences and Health Policy, Wake Forest University School of Medicine, Medical Center Boulevard, Winston-Salem, NC 27157, USA; Department of Psychiatry and Behavioral Sciences, Sylvester Comprehensive Cancer Center, University of Miami Miller School of Medicine, Miami, FL 33136, University of Miami; Dr. John T. Macdonald Foundation Department of Human Genetics, University of Miami Miller School of Medicine, 1501 NW 10th Avenue, Miami, FL 33136, USA; Neuroscience and Ageing Research Unit, Institute for Advanced Medical Research & Training, College of Medicine, University of Ibadan, Ibadan 200212, Nigeria; Department of Medicine, Kwame Nkrumah University of Science & Technology, Private Mail Bag, University Post Office, Kumasi, Ghana; Department of Medicine, University of Ghana Medical School/Korle Bu Teaching Hospital, P.O. Box GP 4236, Accra, Ghana; Gertrude H. Sergievsky Center, Taub Institute for Research on the Aging Brain, Departments of Neurology, Psychiatry, and Epidemiology, College of Physicians and Surgeons, Columbia University, 630 W 168th St, New York, NY 10032, USA; Department of Population & Quantitative Health Sciences, School of Medicine, Case Western Reserve University, 10900 Euclid Ave, Cleveland, OH 44106, USA

**Keywords:** African American, ABCA7, Frameshift Deletion, Lipid Droplet, Alzheimer’s Disease

## Abstract

**INTRODUCTION:** *ABCA7* (*ATP-binding cassette sub-family A member 7*) encodes a lipid transporter linked to Alzheimer’s disease (AD). While common variants confer modest risk in Europeans, a 44-base pair deletion (rs142076058; p.Arg578Alafs) is a strong risk factor in African Americans (AA). Despite this, the biological consequences of this ancestry-specific variant are not well understood.

**METHODS:** We expressed the truncated ABCA7 protein in HEK and HepG2 cells to assess localization and lipid metabolism. Additionally, induced pluripotent stem cell (iPSC)-derived neurons carrying the deletion were compared with isogenic controls.

**RESULTS:** The truncated ABCA7 localized to the plasma membrane similarly to wild type but induced significant lipid droplet accumulation in HepG2 cells and iPSC-derived neurons.

**DISCUSSION:** These findings show that the AA-specific ABCA7 deletion disrupts lipid regulation despite normal localization, suggesting a mechanistic link between impaired lipid homeostasis and increased AD risk. This work underscores the importance of ancestry-specific studies in AD research.

**Highlights:**

- Truncated ABCA7 protein remains stable and correctly localizes to the plasma membrane in HEK293T cells.
- Truncated ABCA7 disrupts lipid droplet regulation in HepG2 cells.
- ABCA7 shows the highest expression in neurons among brain cell types.
- ABCA7 truncation impairs lipid metabolism in neurons.

## 1. BACKGROUND

Alzheimer’s disease (AD) is the most common form of dementia among the elderly, representing about 65% of all dementia cases (1). Pathologically, AD is characterized by extracellular amyloid-β (Aβ) plaques and intracellular neurofibrillary tangles (NFTs) (2, 3). While age is the strongest risk factor, decades of research demonstrate that genetic factors also play a critical role (4-7). Studies aimed at identifying common genetic risk factors for Alzheimer’s disease (AD) have identified 85 loci harboring common genetic variation associated with AD. The first and most well-established genetic risk factor was the ε4 allele of *apolipoprotein E* (*APOE*) (8), which is the strongest known genetic determinant of AD. *APOE* is essential for lipid transport and metabolism in the brain, processes that maintain neuronal function and integrity. Disrupted lipid transport impairs membrane repair, synaptic signaling, and promotes Aβ accumulation. However, the effect of *APOE* ε4 varies across populations, highlighting the importance of ancestry-specific genetic studies.

African American individuals are nearly twice as likely to develop AD compared to individuals of European ancestry, yet APOE ε4 shows weaker or inconsistent associations in African and sub-Saharan African populations. This suggests that additional ancestry-specific risk factors may influence disease susceptibility. One such factor is *ATP-binding cassette sub-family A member 7 (ABCA7)*, a gene implicated in lipid homeostasis, phagocytosis, and Aβ clearance. While common *ABCA7* variants are associated with AD in European ancestry, rare and population-specific variants show stronger effects in African descent populations.

Unique to African ancestry, we identified a 44-base pair (bp) deletion (rs142076058) in *ABCA7* that is significantly associated with AD (9). This *ABCA7* deletion produces a frameshift mutation (p.Arg578Alafs) predicted to encode a truncated protein containing only 2 of the 11 transmembrane domains and lacking both nucleotide-binding domains (9) (Figure 1A).

**Figure 1.**
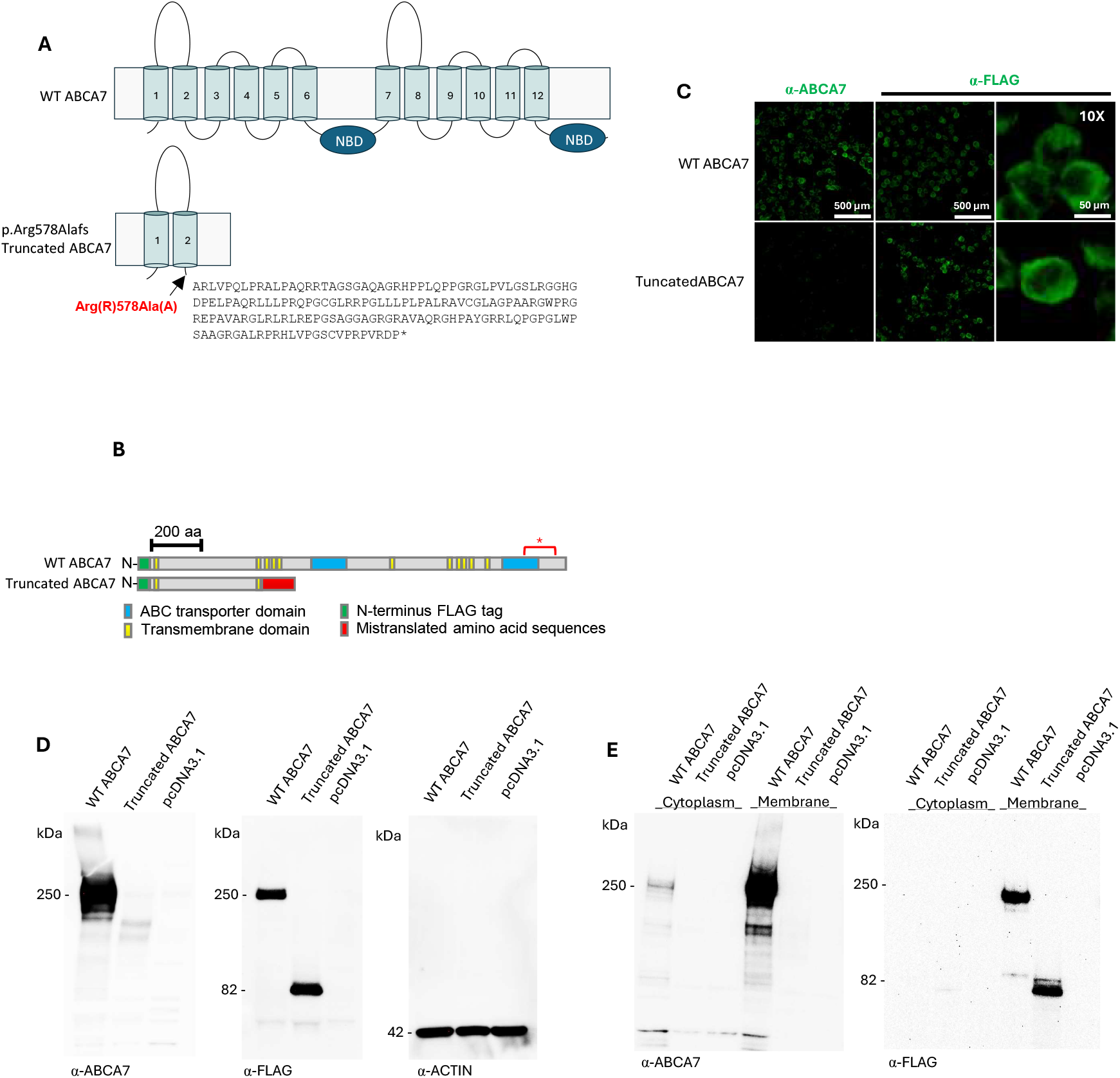
Truncated ABCA7 is stable and localizes at the plasma membrane. (A) Scheme of WT and the predicted truncated ABCA7 proteins at the plasma membrane. WT ABCA7 has 12 hydrophobic transmembrane domains (TMDs) and two nucleotide-binding domains (NBDs). The deletion is predicted to result in an Arg578Alafs^*^171 alteration, which would truncate >70% of the protein and add 171 incorrect amino acids. The predicted truncated ABCA7 contains the first two TMDs without any NBDs. The mistranslated amino sequences are predicted to have no function. (B) A schematic diagram of the N-terminal FLAG-tagged WT ABCA7 and truncated ABCA7. The red star indicates the localization of the epitope for commercially available ABCA7 antibodies. (C) ICC of HEK cells transfected with either WT ABCA7 or truncated ABCA7 probed with anti-FLAG (left panel) or anti-ABCA7 antibodies (middle and right panels). The right panels are magnifications of the corresponding middle panels. (D) Western blot analysis of transfected HEK cells with either anti-ABCA7, anti-FLAG, or anti-Actin as loading control. (E) Western blot analysis of transfected HEK cells subjected to subcellular fractionation with either anti-ABCA7 or anti-FLAG.

Despite its strong genetic association with AD in African ancestry populations, the functional impact of this deletion remains poorly understood. Several factors make this mutation an excellent candidate to study the role of *ABCA7* in increasing the risk for AD. First, our previous demonstration that the mutated mRNA escapes nonsense-mediated decay indicates that the variant likely produces a truncated protein rather than resulting in complete loss of transcript. This suggests that its pathogenicity is more likely due to loss of normal ABCA7 function or gain of function rather than haploinsufficiency due to transcript degradation. Second, our recent genetic interaction studies demonstrate that *APOE 3/4* carriers who also harbor the *ABCA7* 44-bp deletion exhibit a significantly earlier age of AD onset compared to ε3/ε4 carriers without the deletion (10). This finding suggests a potential synergistic interaction between *APOE* ε4 and the *ABCA7* deletion in accelerating disease progression. Given ABCA7’s established role in lipid transport, endolysosomal function, and amyloid-beta clearance, the presence of a truncated protein resulting from this deletion could plausibly amplify AD-related pathology through disrupted homeostatic processes.

Third, a subset of deletion carriers does not go on to develop AD, indicating the potential involvement of genetic modifiers, environmental influences, or compensatory biological mechanisms. These observations underscore the need to define the molecular and cellular impact of the *ABCA7* deletion more precisely. Therefore, we conducted functional studies to investigate the molecular and cellular consequences of the African-specific 44-bp *ABCA7* deletion. These experiments aim to clarify how this variant disrupts ABCA7 function and contributes to AD pathogenesis, ultimately improving our understanding of ancestry-specific genetic risk and guiding the development of more equitable precision medicine strategies.

We demonstrate that the truncated ABCA7 protein is stable and exerts measurable effects on cellular lipid metabolism, supporting a mechanism that may disrupt key processes involved in AD pathogenesis. These findings offer novel insight into the functional role of ABCA7 in the context of ancestry-specific genetic variation and highlight new potential avenues for therapeutic intervention in AD, particularly in populations historically underrepresented in genetic and translational research.

## 2. METHODS

### 2.1 ABCA7 FLAG-tagged plasmid construction

A wild-type *ABCA7* plasmid construct with the human transcript NM_019112 and FLAG-tagged on the N-terminus (EX-A3083-M11) was obtained from GeneCopoeia (Rockville). To mimic the 44-bp deletion (rs142076058), a 308 base pair DNA fragment spanning the deletion was PCR amplified from a custom-made PUC57 plasmid (44-bpDel PUC57 Plasmid – Cat No: SC1691, GenScript). The 44-bp Del pUC57 plasmid was digested with FseI and BstEII (New England Biolabs), and the smaller, 296-base pair fragment including the region with the deletion was purified from a 0.8% agarose gel using a QIAquick gel extraction kit (Qiagen). The GeneCopoeia plasmid was also digested with FseI and BstEII and treated with calf intestinal alkaline phosphatase (CIP, New England Biolabs) for 30 min at 37 ^°^C. The fragment with the deletion was then ligated into the GeneCopoeia backbone overnight with T4 DNA ligase. Sanger sequencing with the following primers (forward: CTTCGTGTACCTGCAAGACC, reverse: GCTGAGCAGGAAGCTCTG) confirmed that the 44-base pair deletion was successfully included in the GeneCopoeia FLAG-tagged plasmid using our previously published methods (9).

### 2.2 Immortalized Cell Lines, Cell Culture, Transfection

HEK/APPsw cells were a gift from Dr. Peter St. George-Hyslop, and HepG2 cells were purchased from ATCC (Cat#: HB-8065). Cell lines were routinely maintained in Dulbecco’s Modified Eagle Medium/Nutrient Mixture F-12 (DMEM/F12, Cat#: 21331020, Gibco), supplemented with 10% fetal bovine serum (FBS, Cat#: A5670701, Gibco) and 1% Penicillin-Streptomycin (P/S, Cat#: 15140122, Gibco) at 37°C in a humidified 5% CO2 atmosphere. PCDNA3.1, ABCA7 WT, or truncated ABCA7 plasmids were transfected into HEK/APPsw and HepG2 cells using the JetPRIME® reagent (Cat#: 89129, Avantor) according to the manufacturer’s protocol. Briefly, cells were seeded 24 hours before transfection. At approximately 70% confluency, the transfection solution containing DNA was added to the cells dropwise. The plate was centrifuged at 180 × g for 5 minutes at room temperature (RT) and incubated for 4 hours before replacing it with fresh DMEM/F12 medium. The transfection with these vectors resulted in high transfection efficiency (>90%). The cells were incubated overnight for post-transfection treatments.

### 2.3 Western Blotting (WB)

PBS-washed cells were lysed with ice-cold RIPA buffer (Cat#: 89900, ThermoFisher Scientific) containing 1X protease inhibitor cocktail (Cat#: P8340, Sigma-Aldrich) on ice for 15 minutes. The lysates were then centrifuged at 13,000 rpm for 10 minutes at 4°C to separate the protein (supernatant) from cellular debris (pellet). The protein concentration of the supernatant was measured using a BCA protein assay kit (Cat#: 23225, ThermoFisher Scientific). Fifteen micrograms of BCA-measured cell lysates were resolved using SDS-PAGE gel electrophoresis on 3%-8% gradient Tris-acetate gels (Cat#: EA03752BOX, ThermoFisher Scientific) and blotted onto 0.45 µm PVDF membrane (Cat#: LC2005, ThermoFisher Scientific) using the Trans-Blot Turbo Transfer System. Membranes were blocked with a 5% final concentration of non-fat dry milk powder for 1 hour. The membranes were probed with primary antibodies, including anti-ABCA7 (Cat#: SC377335, SantaCruz), anti-Flag (Cat#: F1804, Sigma-Aldrich), and anti-Actin (Cat#: AB179467, abcam) overnight at 4°C, followed by probing with secondary antibodies, including anti-mouse and anti-rabbit IgG1 HRP for 1 hour.

### 2.4 Cellular Fractionation into the Plasma Membrane and Cytoplasm

Subcellular fractionation was performed using a Subcellular Protein Fractionation Kit for Cultured Cells (Cat# 78840, ThermoFisher Scientific) according to the manufacturer’s protocol. Briefly, transfected HEK cells (2 × 10^6 cells) were treated with cytoplasmic, membrane, and nuclear extraction buffers containing a protease inhibitor (included in the kit). In each step, subcellular components were collected by centrifugation and used for WB analysis.

### 2.5 Culture and Generation of Human Isogenic iPSCs

Isogenic induced pluripotent stem cell (iPSC) lines containing specific knock-in variants were generated using CRISPR/Cas9-mediated homology-directed repair. Single guide RNAs (sgRNAs) were synthesized by Synthego, reconstituted in nuclease-free 1× TE buffer at 100 µM, pulse vortexed, and incubated on ice for 30–60 minutes before aliquoting and storing at –80°C. To form ribonucleoprotein (RNP) complexes, 140 pmol of sgRNA was combined with 40 pmol of Cas9 nuclease (EnGen Spy Cas9 NLS, Cat# M0646T, New England Biolabs) in Lonza P3 Nucleofector solution (Cat# V4XP-3032) at a 3.5:1 molar ratio, then incubated at room temperature for 10 minutes. iPSCs were cultured in mTeSR Plus medium (StemCell Technologies) on vitronectin-coated plates (Cat# A14700, ThermoFisher Scientific), and pre-treated for at least one hour with 10 µM Y27632 and 2 µM thiazovivin to improve cell viability. After dissociation with Accutase, cells were washed in DMEM/F12 (Cat# 10565042, ThermoFisher Scientific) and resuspended in nucleofection solution containing 3 µM donor single-stranded oligodeoxynucleotide (ssODN) (IDT Ultramer, 100 µM stock). A total of 2 × 10□ cells were mixed with the RNP complexes and electroporated using the 4D-Nucleofector X Unit (Lonza) with program CA137. Transfected cells were plated onto vitronectin-coated 12-well plates and cultured in mTeSR Plus supplemented with 10 µM Y27632 and 2 µM thiazovivin. Following recovery, bulk-transfected cells were cryopreserved, and genomic DNA was extracted from cell pellets using QuickExtract DNA Extraction Solution (Cat# QE09050, Qiagen). Editing efficiency was evaluated through Sanger sequencing and analyzed with the ICE tool (Synthego). To isolate single-cell clones, limiting dilution was performed at 1 cell per 100 µL in mTeSR Plus with CloneR2 (StemCell Technologies), plated onto rhLaminin-521–coated 96-well plates (Cat# A29248, ThermoFisher Scientific). After 7–10 days, emerging colonies were visually screened, cryopreserved in CryoStor CS10 (StemCell Technologies), and genotyped. Selected clones were thawed into mTeSR Plus supplemented with CloneR2 (Cat#: 100-0691, StemCell), expanded, and verified by Sanger sequencing to confirm successful KI incorporation.

During the generation of CRISPR-edited lines, efforts to obtain homozygous or heterozygous clones carrying a 44-bp deletion in ABCA7 were challenging, yielding either no clones or a very small number (<3%), none of which survived the iPSC expansion process (Supplemental Figure 1a). However, a higher number of clones (>10%) with a 47-bp homozygous or heterozygous deletion in ABCA7 were successfully obtained. Since the 47-bp deletion removed an additional single codon (Leu(L)577Ala(A)), minimally altering the resulting protein compared to the 44-bp deletion (Arg(R)578Ala(A)) (Supplementary Figure 1B&C), the 47-bp deletion clones were selected as functional equivalents of the 44-bp deletion in ABCA7. The generated iPSC lines were fully validated for pluripotency and genomic stability using G-banding karyotyping (Supplemental Figure 2). Additionally, Sanger sequencing was performed to confirm the genotypes and assess potential CRISPR off-target effects in the ABCA7 deletion lines. CRISPR-generated isogenic ABCA7 deletion iPSC lines were differentiated into neurons by a conventional neuronal induction approach by a transdifferentiation approach to overexpress

*Neurogenin-2* (*NGN2*) using home-brew cocktails. Throughout neuronal induction and maturation, no significant developmental defects were observed in neuronal structures (Supplemental Figure 3).

### 2.6 Transdifferentiation to iPSC-Derived Neurons (iN)

PB-Ngn2-EF1a-Puro-BFP and PB-transpose plasmids were gifted by Dr. Samuele Marro from the Nash Family Department of Neuroscience and the Black Family Stem Cell Institute at the Icahn School of Medicine at Mount Sinai. The generation of iN cells from human iPSCs was described previously (11, 12). Briefly, iPSC cultures at 70-80% confluency were visually inspected to ensure proper morphology and the absence of untargeted differentiation. On day 0, 3 × 10^4 single-cell dissociated iPSCs were seeded onto a 1w/6w plate in mTeSR+ (Cat#: 100-0276, StemCell). On day 1, cells were transfected with PB-Ngn2-EF1a-Puro-BFP and PB-transpose plasmids using Lipofectamine Stem Transfection Reagent (Cat#: STEM00015, ThermoFisher Scientific) according to the manufacturer’s protocol. After 4 hours of transfection, 2 mL of mTeSR+ was added, and the cells were incubated overnight. On day 2, the media was aspirated and replaced with fresh 1.5 mL mTeSR+. On days 3 and 5, transfection selection was initiated by adding puromycin (final concentration of 2 µg/mL, Cat#: 540411, Sigma-Aldrich). On day 6, cells were passed onto a 6w/6w plate for full puromycin selection. On days 7 and 9, cells were continued on mTeSR+ with puromycin. By days 11-13, when the confluency reached over 70%, integration of PB-Ngn2-EF1a-Puro-BFP was confirmed (>95% BFP expression) and neuronal induction and maturation were initiated. On day 0, 1 × 10^6 single-cell dissociated cells were seeded onto a Matrigel-coated 1w/6w plate in a home-brewed neuronal induction media (11) containing 2 µg/mL doxycycline with CultureCEPT (Cat#: A56799, ThermoFisher Scientific). On days 1 and 2, cells were washed and replaced with induction media without CultureCEPT. On day 3, cells were treated with 1 µM uridine (Cat#: U3003, Sigma-Aldrich) and fluorodeoxyuridine (Cat#: F0503, Sigma-Aldrich) to suppress mitotic events. On day 4, cells were replated (1-2 × 10^6 cells) onto a PLO-coated 1w/6w plate in home-brewed neuronal maturation media I (11) with CultureCEPT. On day 5, the media was replaced without CultureCEPT. On day 8, cells were fed with a 1/2 media change. On days 11, 13, 17, 23, and 26, cells were fed with home-brewed neuronal maturation media II (11). On day 28, mature iNs were harvested for desired assays.

### 2.7 Immunocytochemistry

Cells were fixed in 4% methanol-free formaldehyde (Image-iT™ Fixative Solutions, Cat#: I28800, ThermoFisher Scientific) at RT for 15 minutes, washed three times with PBS, then blocked/permeabilized with 10% Normal Donkey Serum (Cat#: 017-000-121, Jackson ImmunoResearch), 0.25% Triton X-100, and 0.05% Tween-20 for 1 hour. Cells were then incubated with primary antibodies, including anti-ABCA7 (Cat#: SC377335, SantaCruz), anti-Flag (Cat#: F1804, Sigma-Aldrich), β-Tub III (Cat#: AB78078, abcam), and MAP2 (Cat#: 13-1500, ThermoFisher Scientific), in a blocking solution overnight at 4°C or for 1 hour at RT. Cells were washed three times with PBS, then incubated with secondary antibodies in a blocking buffer for 1 hour at RT. Cells were incubated with Hoechst 33342 (Cat#: R37605, ThermoFisher Scientific) for 20 minutes at RT, followed by 3X PBS washes for 5 minutes. Cells were washed once more and replenished with PBS for imaging.

### 2.8 Treatments and Lipid Droplet Assay

Transfected HepG2 cells with *ABCA7* WT, *ABCA7* Del, or empty pcDNA3.1 plasmids were treated with either 500 µM BSA-Oleic acid (OA, Cat#: 29557, Cayman) to induce lipid droplet accumulation or control BSA (Cat#: 34932, Cayman) overnight. LDs were visualized using LipidSpot™ 610 dye (70069, Biotium) for 30 minutes in live cells, followed by fixation and subsequent immunofluorescence staining. Only cells that showed a positive signal for expression of the recombinant version of ABCA7 (as determined by immunocytochemical staining for the N-terminal FLAG tag) were assessed for the determination of lipid droplet formation.

### 2.9 Light Microscopy Image Acquisition and Analysis

Images were acquired using an automated high-resolution fluorescence microscope (BZ-X800, Keyence) equipped with DAPI, Green, Texas Red, and FarRed channels. Acquisition settings, such as exposure and image quality of individual fluorescence channels, were kept identical to avoid biases across imaging. Z-planes of 0.2 µm thickness were acquired. To quantify the abundance of LD in each cell, images of representative cells were initially cropped into a 1-inch square. Using ImageJ, the border of an individual cell was manually outlined, and the images were converted to 8-bit for thresholding. Signals found outside the cell border were manually excluded using the paintbrush tool. Thresholding was performed to measure the percentage of lipid area in each cell.

### 2.10 Statistical analysis

All statistical analyses were performed in JMP Pro 17 (JMP Version 17, SAS Institute Inc.). Two-tailed Student’s t-test was used to compare between groups. A two-tailed p < 0.05 was considered significant for all statistical tests. The post hoc power analysis was conducted by using Tukey’s post hoc analysis.

## 3. RESULTS

### 3.1 The truncated ABCA7 protein is stable and is localized to the plasma membrane in HEK293T cells

To investigate whether the 44-bp *ABCA7* deletion affects protein stability and localization, we transfected HEK cells with a plasmid encoding either the N-terminally tagged full-length (WT ABCA7) or the truncated ABCA7 predicted to result from the 44-bp deletion (Figure 1B). Immunocytochemistry (ICC) analysis for FLAG immunodetection showed that both the full-length and truncated ABCA7 were localized at the cell membrane (Figure 1C). As expected, an antibody against the C-terminal domain of ABCA7 detected WT ABCA7 but not the truncated ABCA7 protein. This finding suggested that the truncated ABCA7 protein was stable and correctly localized.

These results were validated by western blot analysis (Figure 1D). Immunoblotting using a FLAG antibody showed similar levels of WT ABCA7 and truncated ABCA7, with each migrating based on its expected sizes (approximately 240 kDa for WT and 82 kDa for the truncated version). Additionally, subcellular fractionation was performed to separate the cytoplasmic and plasma membrane fractions from the transfected cells (Figure 1E). Both WT ABCA7 and truncated ABCA7 localized to the plasma membrane, confirming the results of the immunocytochemical analysis (Figure 1C). Together, these findings indicate that the truncated ABCA7 protein is stable, correctly transported, and localized to the plasma membrane in transfected HEK cells.

### 3.2 The truncated ABCA7 protein affects lipid droplet formation in HepG2 cells

To test if truncated ABCA7 could influence lipid droplet (LD) accumulation, we transfected HepG2 cells either with WT or truncated ABCA7 and compared LD accumulation by staining with LipidSpot. We utilized the liver-derived HepG2 cells, which exhibit lipid droplet formation when exposed to fatty acids like oleic and palmitic acid (13). The results showed minimal lipid droplet accumulation in HepG2 cells exposed to BSA (vehicle control), and this was not affected by overexpression of any of the *ABCA7* versions (Figure 2A). As expected, Oleic Acid (OA) exposure led to a pronounced increase in LD accumulation across transfected HepG2 cells. Notably, in these conditions, overexpression of WT ABCA7 significantly reduced lipid droplet accumulation compared to the empty vector control (pcDNA3.1), whereas cells overexpressing the truncated ABCA7 were not significantly different from the empty vector control and were significantly higher than those observed in WT ABCA7-transfected cells (Figure 2B). Furthermore, the truncated ABCA7 exhibited a trend toward increased lipid droplet accumulation compared to the empty vector control. These findings suggest that WT ABCA7 plays a role in regulating lipid droplet accumulation in HepG2 cells, a function that seems affected in the truncated ABCA7.

**Figure 2.**
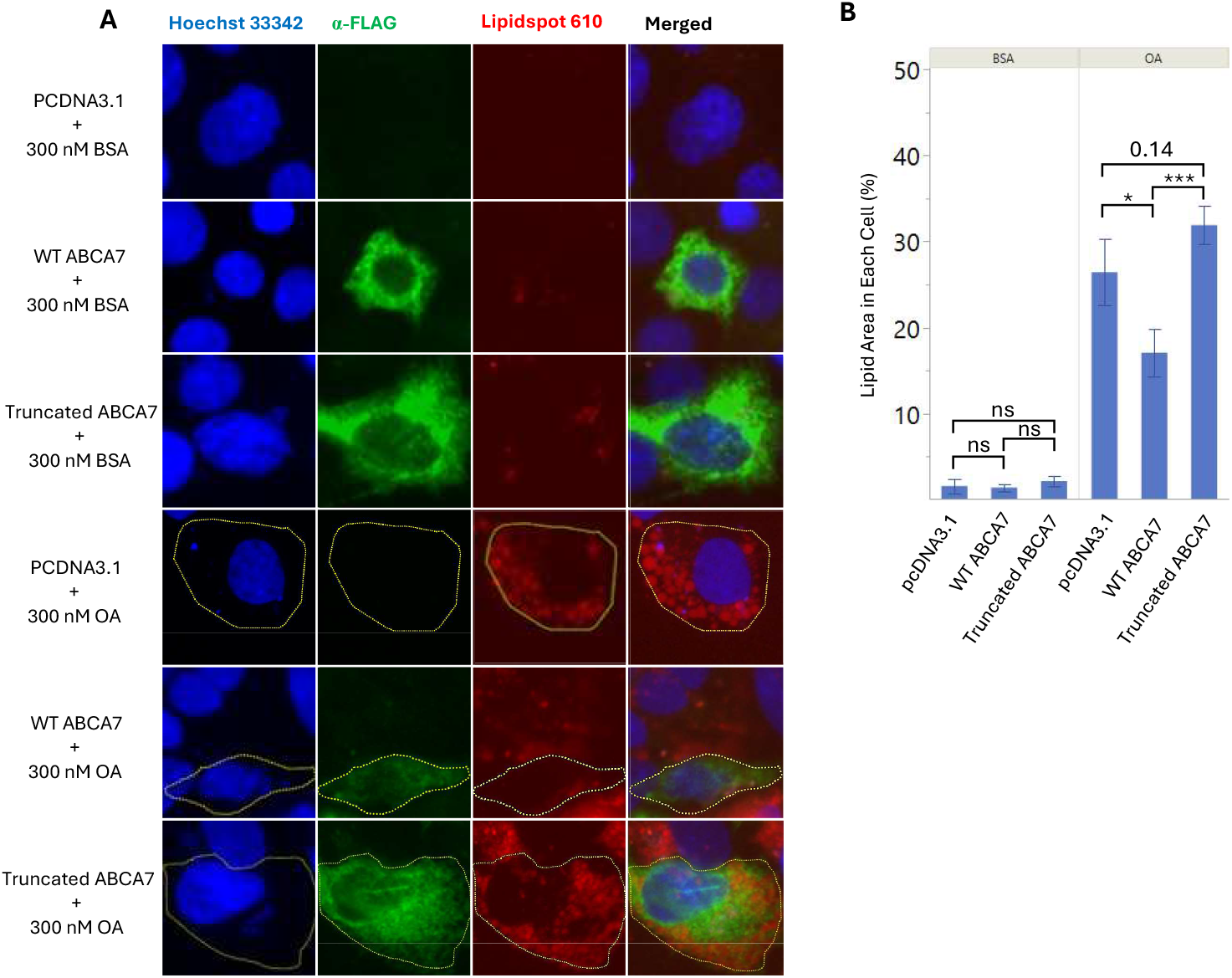
Functional effect in lipid droplet metabolism of truncated ABCA7 in transfected HepG2 cells. (A) Images of HepG2 cells transfected with empty vector, WT ABCA7, or truncated ABCA7 stained for the detection of FLAG and lipid droplets in the presence of 300 nM BSA (top three panels) or OA (bottom three panels). (B) Quantitation of lipid droplet area (%) per transfected cell in the presence of 300 nM BSA (left three bars) or OA (right three bars) shows that the effect of the ABCA truncated protein is different than the WT. These experiments were performed twice independently with technical triplicates. Data represent the mean +/- s.e.m., ^*^p<0.05, ^***^p<0.001, ns: non-significant (one-way ANOVA followed by Student’s t post hoc test).

### 3.3 *ABCA7* is expressed at the highest level in neurons

Previous studies have reported that *ABCA7* transcriptional expression is generally low across various brain cell types (14). Our single-nuclei RNA sequencing (snRNAseq) data from human frontal cortex showed that *ABCA7* expression was highest in neurons (Figure 3A) (15). Figure 3 shows a comparison with *APOE* expression levels, known to be minimal in neurons, revealing the extent of expression of *ABCA7*. In addition, we also analyzed *ABCA7* expression levels in induced pluripotent stem cell (iPSC)-derived neural spheroids, which include neurons, astrocytes, and oligodendrocytes (Figure 3B) (16). The analysis was conducted on neural spheroids derived from four iPSC lines representing individuals of African, Amerindian, and European ancestry. Neurons in the iPSC-derived spheroid exhibited the highest *ABCA7* transcript levels.

**Figure 3.**
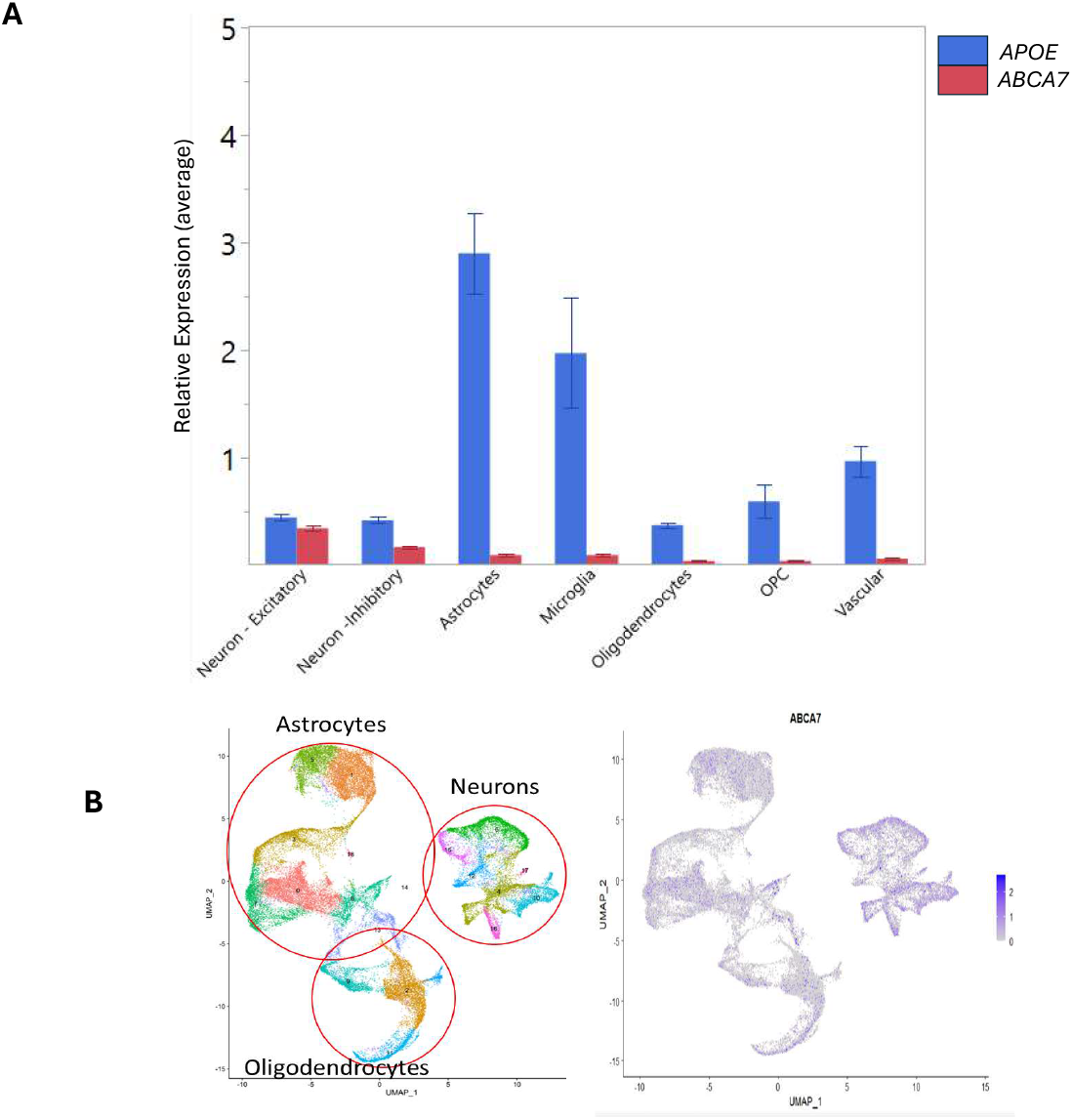
*ABCA7* is expressed brain-wide at relatively low levels, with the highest expression in neurons. (A) *ABCA7* and *APOE* expression level comparison across cell types from single-nucleus RNA sequencing of the frontal cortex of AD patients. (B) UMAP of single-cell RNAseq of iPSC-derived cortical spheroids from 12 individuals, colored based on cell-type cluster (left) or *ABCA7* expression levels (right).

### 3.4 ABCA7 truncation alters lipid metabolism in neurons

The relatively high expression of *ABCA7* in neurons prompted us to analyze the effect of the ABCA7 truncation in this cell type. Thus, we generated isogenic iPSC lines containing either WT *ABCA7* or lines homozygous or heterozygous for the *ABCA7* deletion by CRISPR gene editing. The *ABCA7* deletion was introduced into iPSCs derived from three independent, cognitively unimpaired, non-deletion carrier African American individuals, resulting in three sets of isogenic *ABCA7* lines. Each isogenic set included a CRISPR-unedited control line, a heterozygous *ABCA7* deletion line, and a homozygous *ABCA7* deletion line (Figure 4A). These cells were induced to become human glutamatergic neurons via forced expression of *NGN2* (17).

**Figure 4.**
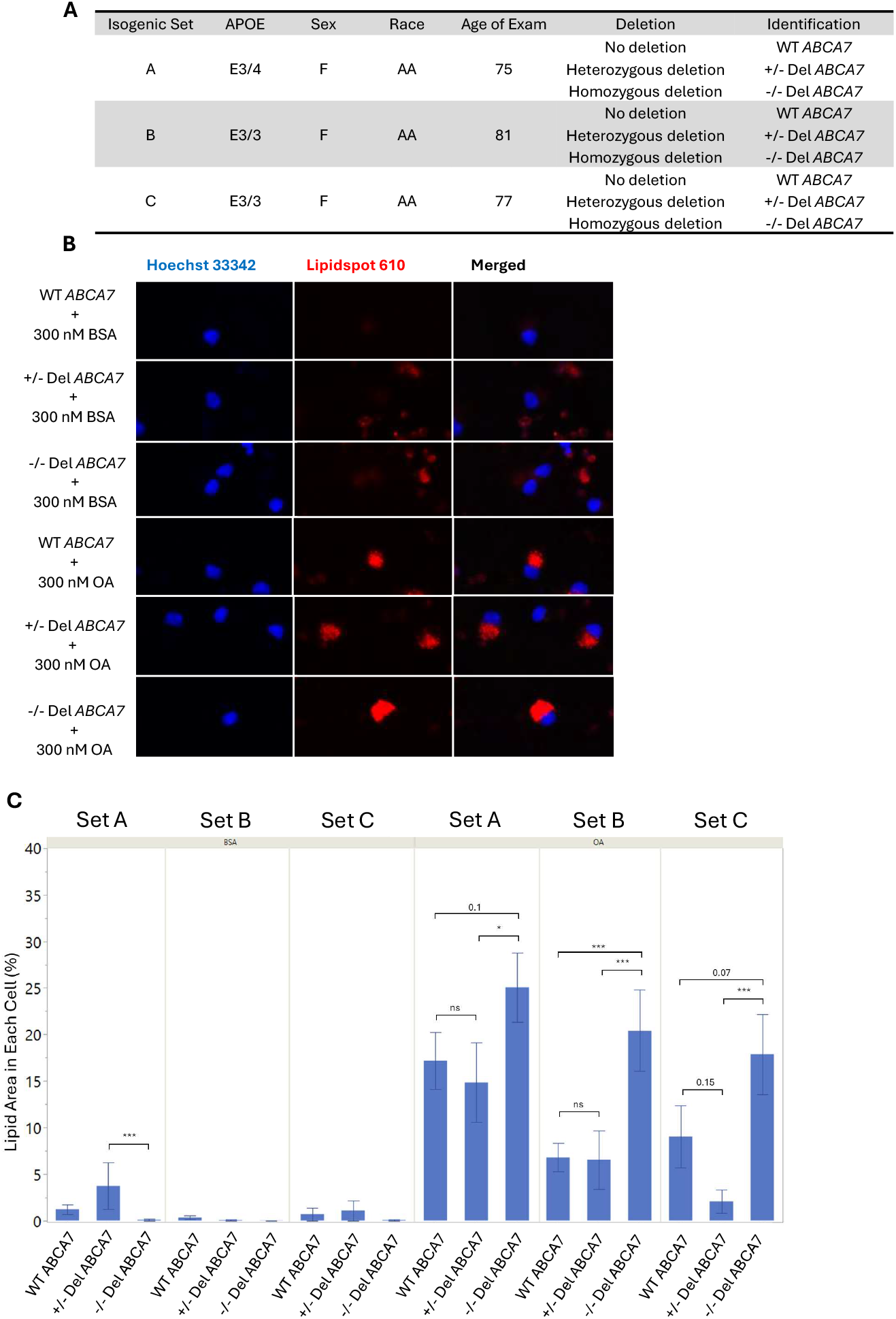
Increased neuronal lipid droplet accumulation in homozygous carriers of the 44-bp deletion in *ABCA7*. (A) Table of isogenic iPSC lines derived from unaffected individuals with an African ancestry (AA) background. CRISPR-based genome editing was used to introduce the AA-specific *ABCA7* deletion in homozygous or heterozygous states. (B) Representative immunocytochemistry (ICC) images showing lipid droplet accumulation in isogenic neurons carrying wild-type (WT), heterozygous *ABCA7* deletion, or homozygous *ABCA7* deletion following treatment with 200 nM BSA (top three panels) or oleic acid (OA, bottom three panels). (C) Quantitation of lipid area (%) in individual isogenic neurons stained with LipidSpot. The lipid area from ten individual neurons per well was averaged. These experiments were performed with technical triplicates. Data are presented as mean ± s.e.m. ^*^p < 0.01, ^***^p < 0.001 (one-way ANOVA followed by Student’s t-test for post hoc analysis).

To test whether the *ABCA7* deletion affected lipid droplet accumulation in neurons, as observed in HepG2 cells, we exposed the nine isogenic ABCA7 neuronal lines to either BSA (vehicle control) or oleic acid (OA) and quantified lipid droplet accumulation (Figure 4B). Lipidspot quantification revealed that neurons exposed to BSA did not exhibit significant lipid droplet accumulation. However, OA exposure induced lipid droplet formation across all isogenic neuronal sets (Figure 4C). Analysis of the three independent isogenic sets (A, B, and C) revealed a consistent pattern, where neurons with homozygous *ABCA7* deletion exhibited a significant increase in lipid droplet accumulation compared to both WT *ABCA7* and heterozygous *ABCA7* deletion neurons (Figure 4D).

## 4. DISCUSSION

This study sheds light on the functional consequences of the African ancestry-enriched 44-bp deletion in *ABCA7*, a variant strongly associated with increased Alzheimer’s disease (AD) risk. Unlike most nonsense mutations in *ABCA7*, which produce truncated proteins that are mislocalized or degraded (18), we find that the truncated ABCA7 protein resulting from the 44-bp deletion is both stable and correctly trafficked to the plasma membrane in HEK cells. This unexpected finding challenges the widely held assumption that AD risk associated with *ABCA7* truncations operates solely through haploinsufficiency and opens the door to alternative pathogenic mechanisms. The presence of a membrane-localized, truncated ABCA7 protein raises the possibility of dominant-negative effects, where the mutant protein may interfere with the wild-type allele’s function in heterozygous carriers. This model aligns with clinical observations that heterozygous and a few homozygous *ABCA7* 44-bp deletion carriers often present with similar disease severity and age of onset (19).

One significant finding from our study is the demonstration that WT ABCA7 suppresses lipid droplet (LD) accumulation, a regulatory function that is lost in cells expressing the truncated ABCA7. This defect observed in liver-derived HepG2 cells was consistent with the observation of significantly elevated LD levels in iPSC-derived human neurons homozygous for the deletion. These results agree with prior work in animal models, including *Drosophila*, showing that *ABCA7* loss leads to lipid accumulation (20), and in mice, where *ABCA7* knockout impairs lipid metabolism and endolysosomal function (18, 21).

LDs are cellular organelles essential for lipid storage, metabolism, signaling, and protection against lipotoxicity (22). The significance of LDs in Alzheimer’s Disease (AD) has been extensively documented. For instance, microglial LD accumulation, associated with pathogenic *APOE* variants, induces Tau phosphorylation and subsequent neurotoxicity (23). Furthermore, APOE is transported to astrocyte lipid droplets, where it influences triglyceride saturation and the size of the droplets (24). Although neurons typically possess a lower intrinsic capacity for LD storage compared to glia (25), maintaining lipid homeostasis is crucial for synaptic health and resistance to metabolic stress, particularly during aging. The observation that *ABCA7* is expressed at the highest levels in the brain in neurons — based on both single-nucleus RNA sequencing of human brain tissue and analysis of iPSC-derived neural spheroids — underscores the potential for a cell-autonomous, neuron-specific role of *ABCA7* in lipid regulation. Our identification of a functional defect caused by the *ABCA7* deletion — a truncated *ABCA7* that fails to replicate the regulatory function of the wild-type protein in controlling lipid droplet accumulation in transfected cells — suggests that this variant may influence AD risk through disrupted lipid processing pathways. This finding was further supported by studies using isogenic human iPSC-derived neurons, where we observed increased LD accumulation in homozygous deletion cells. Our findings align with recent reports demonstrating a critical role for *ABCA7* in neuronal lipid homeostasis (26). These mechanisms may be especially relevant in the context of *APOE4*, which also disrupts lipid handling and dynergizes with ABCA7 dysfunction (27). The convergence of these pathways points toward a lipid-centered model of AD risk and suggests that impaired lipid metabolism in neurons is the pathological mechanism that leads to the increased risk seen in *ABCA7* 44-bp deletion carriers.

An intriguing aspect of our results is that heterozygous deletion neurons did not show a significant lipid phenotype, despite the clinical significance of heterozygous 44-bp deletion carriers. Several factors may account for this discrepancy. The iPSC-derived neurons used here model early developmental stages and may lack aging-associated stressors necessary to reveal subtle deficits. Alternatively, compensatory mechanisms in these *in vitro* systems may buffer against the partial loss of *ABCA7* function. It is also plausible that the 44-bp deletion exhibits a threshold effect, where a full loss of function is required to produce a measurable cellular phenotype in culture. Yet, given the clinical equivalence between heterozygotes and homozygotes, it seems more likely that the *in vitro* model lacks the sensitivity or maturity needed to detect pathogenic changes in heterozygotes that accumulate over time *in vivo*.

Our recent epidemiological studies indicate that the *ABCA7* deletion reduces the age of onset in individuals carrying the *APOE 3/4* allele compared to those without the deletion or with other *APOE* genotypes (10). Indeed, a previous report suggested that ABCA7-APOE interactions may influence memory function (28). APOE plays a crucial role in lipidation and lipid transport for maintaining lipid homeostasis in the brain (29). APOE localizes to the surface of LD in astrocytes, regulating lipid droplet size, turnover, and sensitivity to peroxidation (24). APOE4 has been shown to dysregulate lipid droplet dynamics, potentially contributing to its strong risk for AD (24). Therefore, the deletion in *ABCA7* may further hinder the transport of lipids seen in APOE4 carriers, leading to an earlier age of onset. Further studies investigating the functional interaction between *ABCA7* deletions and *APOE* alleles are necessary to clarify this relationship and could provide greater clarity into the pathological mechanism of APOE4.

## 5. Conclusion

Taken together, these findings support a model in which the 44-bp deletion in *ABCA7* increases AD risk through impaired regulation of lipid homeostasis, with loss of *ABCA7* function leading to LD accumulation in neurons. Whether this occurs via haploinsufficiency, dominant-negative effects, or context-dependent loss of function remains to be fully elucidated. Importantly, our study is the first to demonstrate that the truncated ABCA7 protein resulting from the 44-bp deletion is stable and traffics to the plasma membrane, providing critical mechanistic insight into how this variant may disrupt lipid metabolism.

These results not only lay the groundwork for understanding how ABCA7-mediated lipid dysregulation contributes to AD pathogenesis but also highlight the importance of ancestry-informed functional genomics in uncovering population-specific mechanisms by which *ABCA7* variants drive neurodegeneration.

## Acknowledgements

**Acknowledgements, Conflicts, Funding Sources, and Consent Statement**

## Acknowledgments

We are also grateful to the many participants, researchers, and staff who contributed significantly to this study.

## Consent Statement

No human data was derived and, therefore, consent was not necessary.

## List of Abbreviations

AD: Alzheimer’s Disease
ABCA7: ATP Binding Cassette Subfamily A Member 7
AA: African American
LD: lipid droplet
iPSC: induced pluripotent stem cell
OA: Oleic Acid

