## Supplemental figure 1 for "Disrupted Lipid Homeostasis as a Pathogenic Mechanism in *ABCA7*-Associated Alzheimer’s Disease Risk"

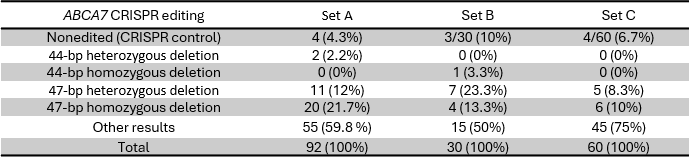

CRISPR control

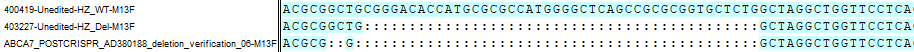

44-bp Deletion

44-bp Deletion *ABCA7*

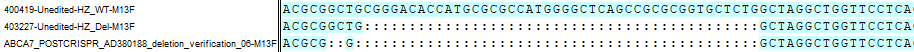

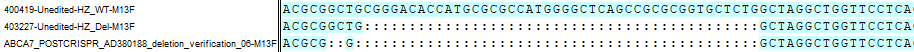

47-bp Deletion

47-bp Deletion *ABCA7*

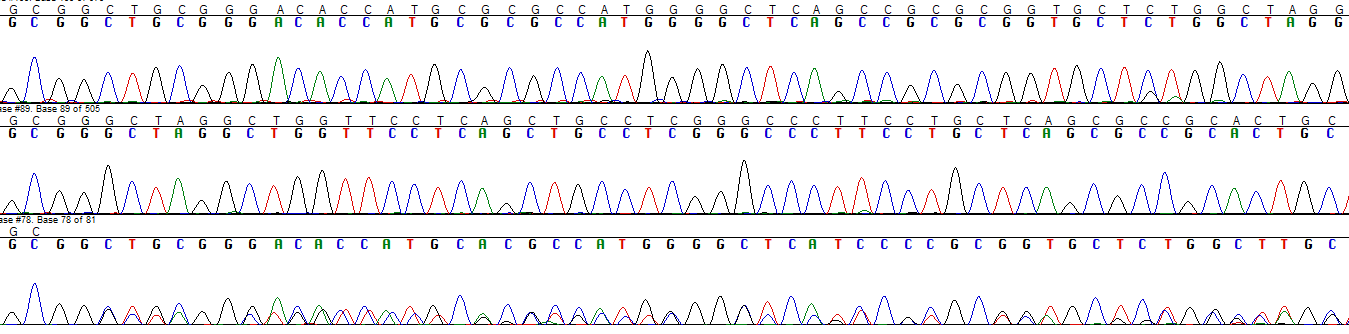

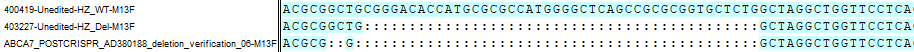

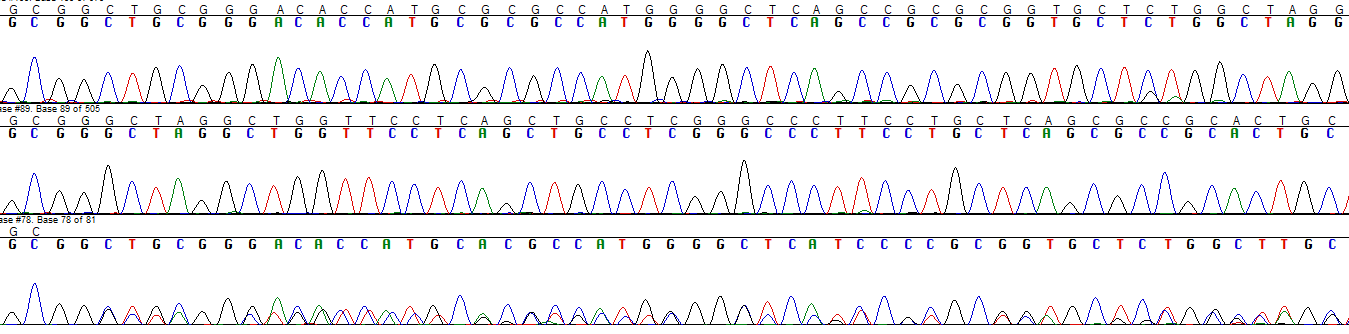

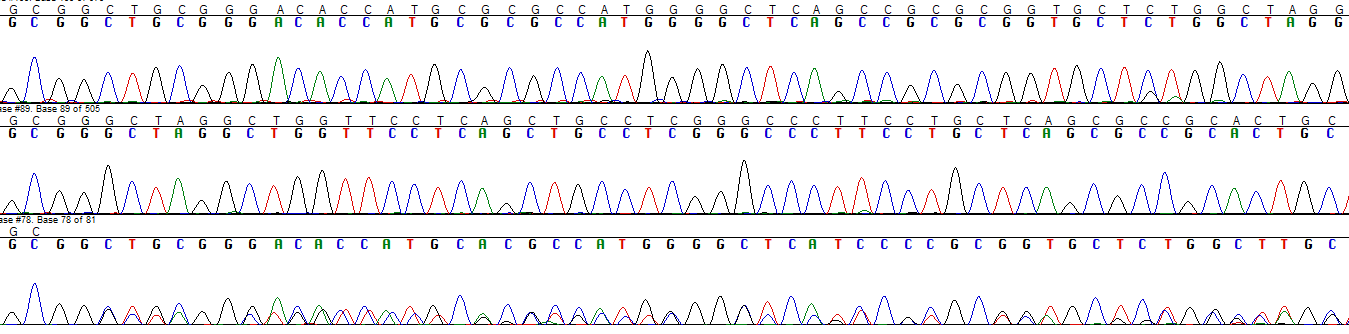

47-bp Homozygous Deletion *ABCA7*

47-bp Heterozygous Deletion *ABCA7*

**B**

**A**

Supplemental Figure 1.

**C**

p.Arg578Alafs

44-bp Del ABCA7

**Arg(R)578Ala(A)**

ARLVPQLPRALPAQRRTAGSGAQAGRHPPLQPPGRGLPVLGSLRGGHGDPELPAQRLLLPRQPGCGLRRPGLLLPLPALRAVCGLAGPAARGWPRGREPAVARGLRLRLREPGSAGGAGRGRAVAQRGHPAYGRRLQPGPGLWPSAAGRGALRPRHLVPGSCVPRPVRDP*

1	2

p.Lue577Alafs

47-bp Del ABCA7

**Leu(L)577Ala(A)**

ARLVPQLPRALPAQRRTAGSGAQAGRHPPLQPPGRGLPVLGSLRGGHGDPELPAQRLLLPRQPGCGLRRPGLLLPLPALRAVCGLAGPAARGWPRGREPAVARGLRLRLREPGSAGGAGRGRAVAQRGHPAYGRRLQPGPGLWPSAAGRGALRPRHLVPGSCVPRPVRDP*

1	2

**Supplemental Figure 1.** Development of CRISPR isogenic ABCA7 iPSC lines and karyotype quality control. (A) CRISPR control, 44-bp heterozygous and homozygous *ABCA7* deletion clones were identified. Less 44-bp *ABCA7* deletion clones were obtained compared to 47-bp *ABCA7* deletion clones. (B) Sanger sequencing chromatogram confirmed the 44 and 47-bp deletion in *ABCA7* using CRISPR-Cas9 gene editing. (C) A schematic protein diagram of the predicted proteins encoded by the 44 and 47-bp deletions at the plasma membrane. The predicted protein from the 47-bp deletion loses another amino acid due to the three additional nucleotides removed, but it maintains the same remaining protein sequences, including the early termination, as the 44-bp *ABCA7* deletion.
