## Supplemental figure 2 for "Disrupted Lipid Homeostasis as a Pathogenic Mechanism in *ABCA7*-Associated Alzheimer’s Disease Risk"

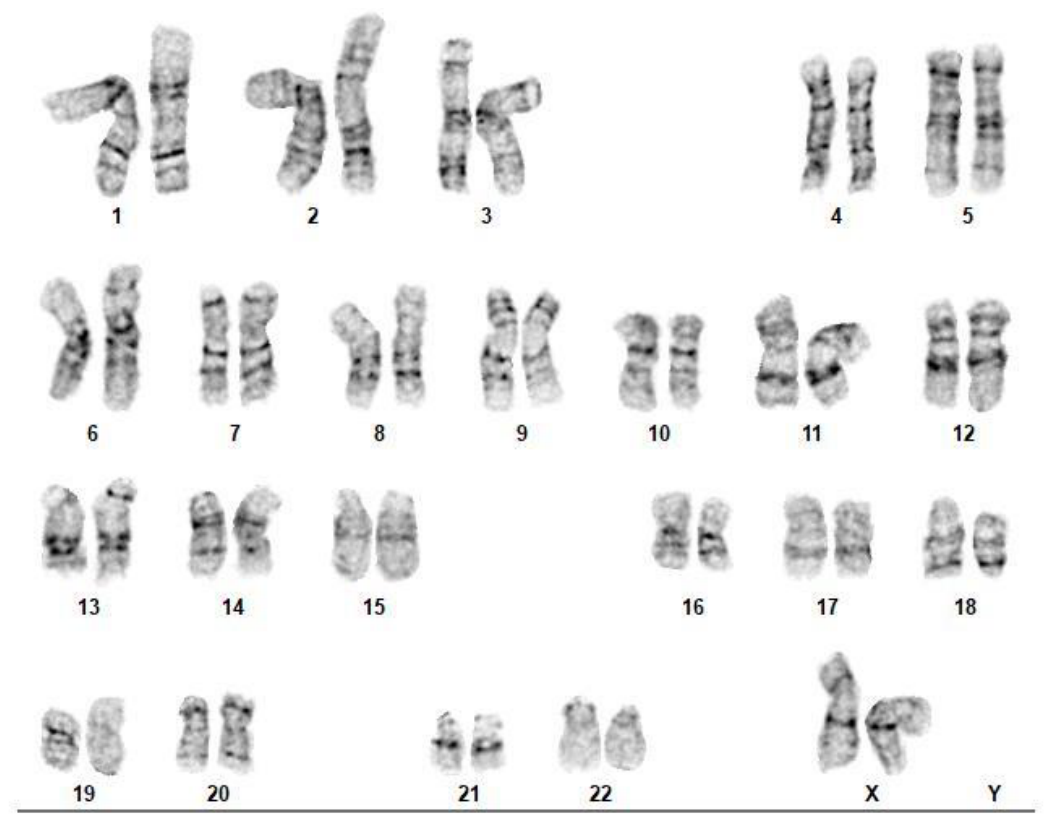

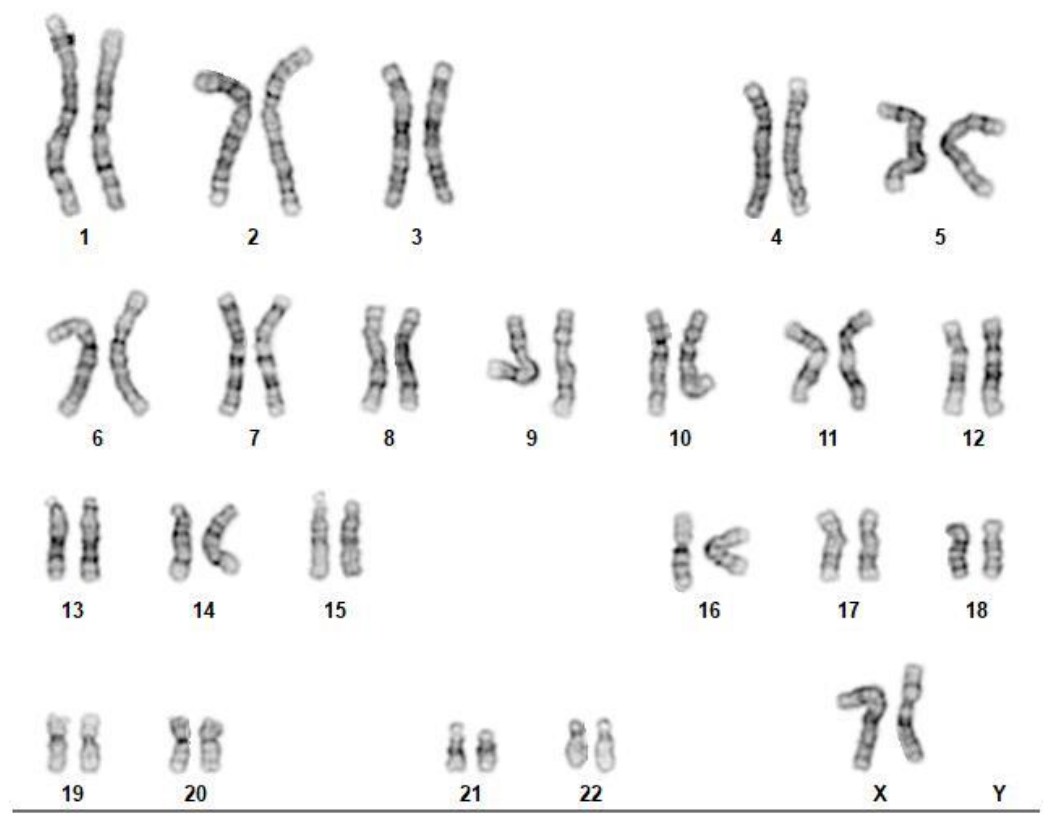

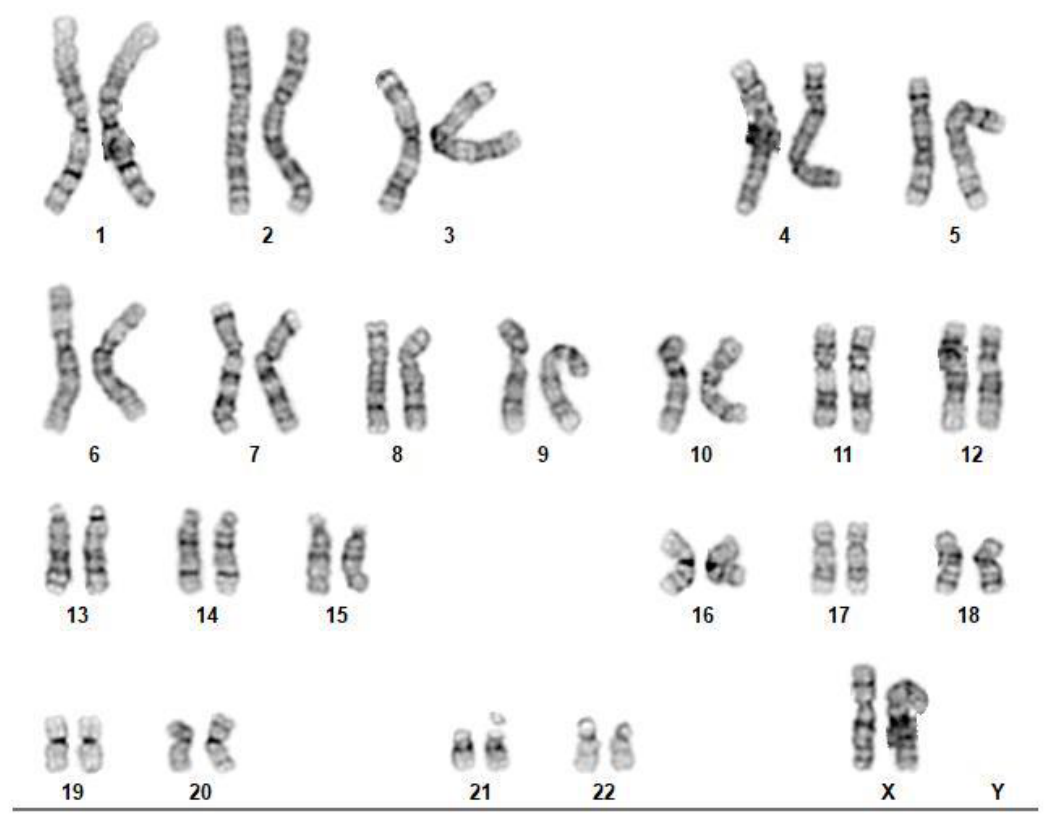

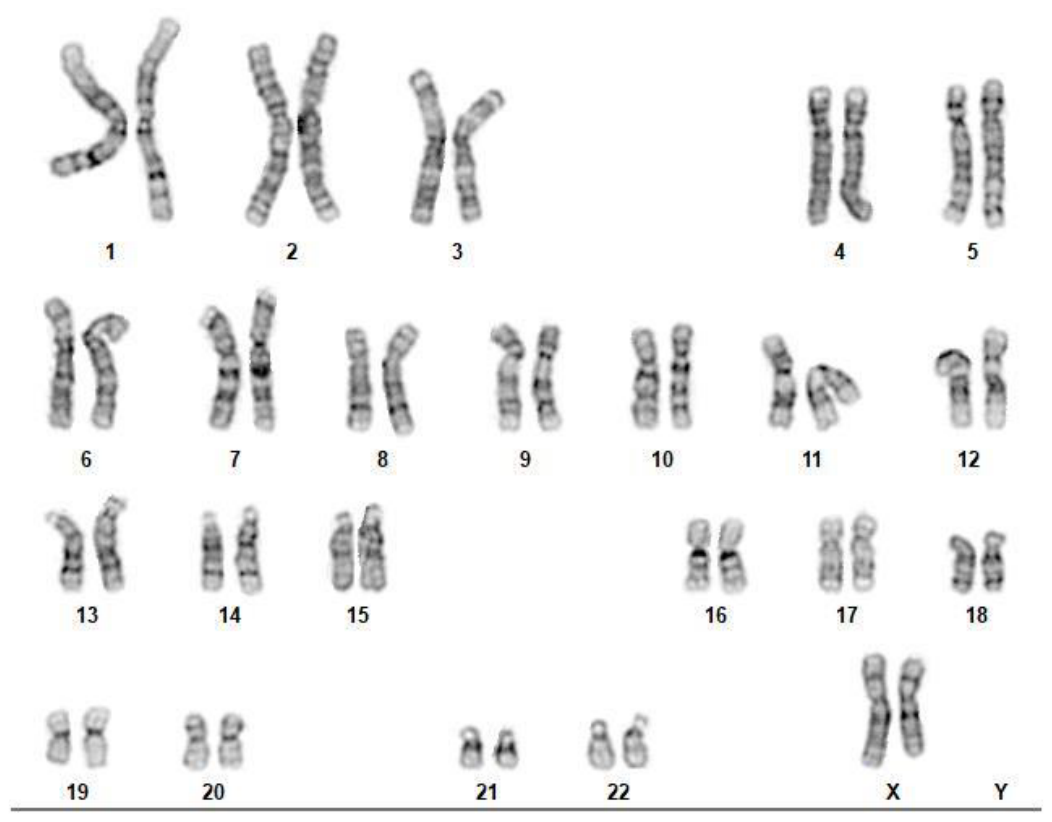

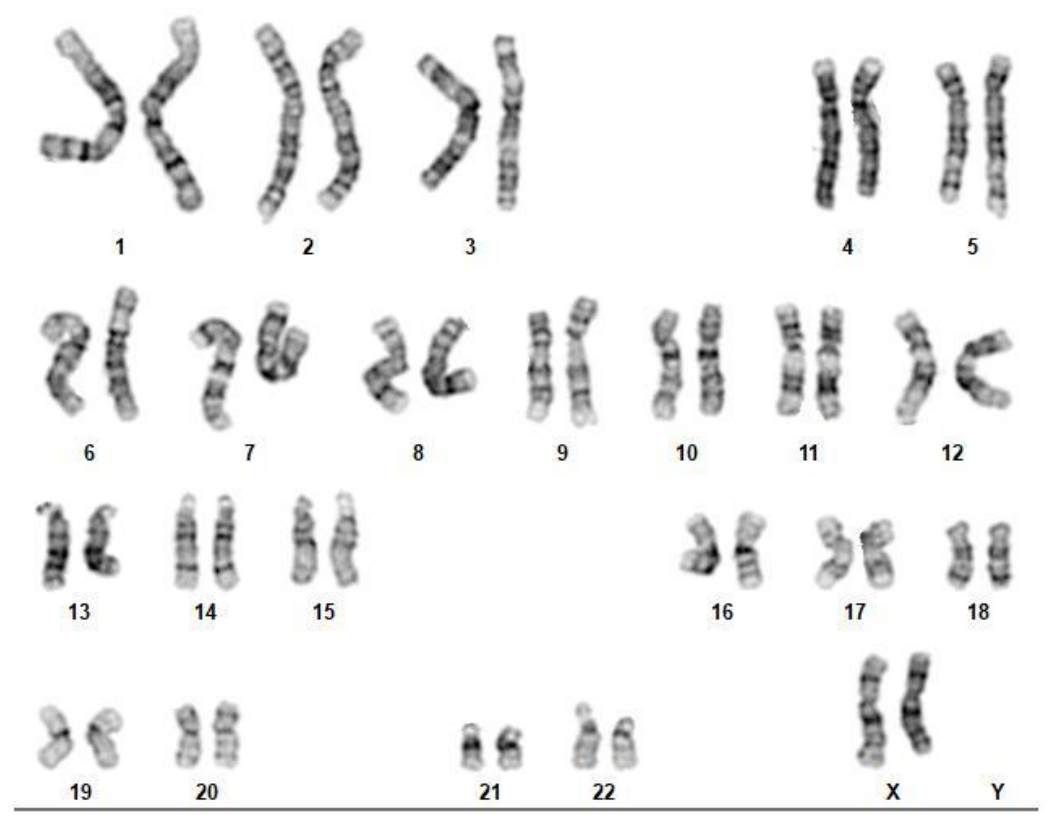

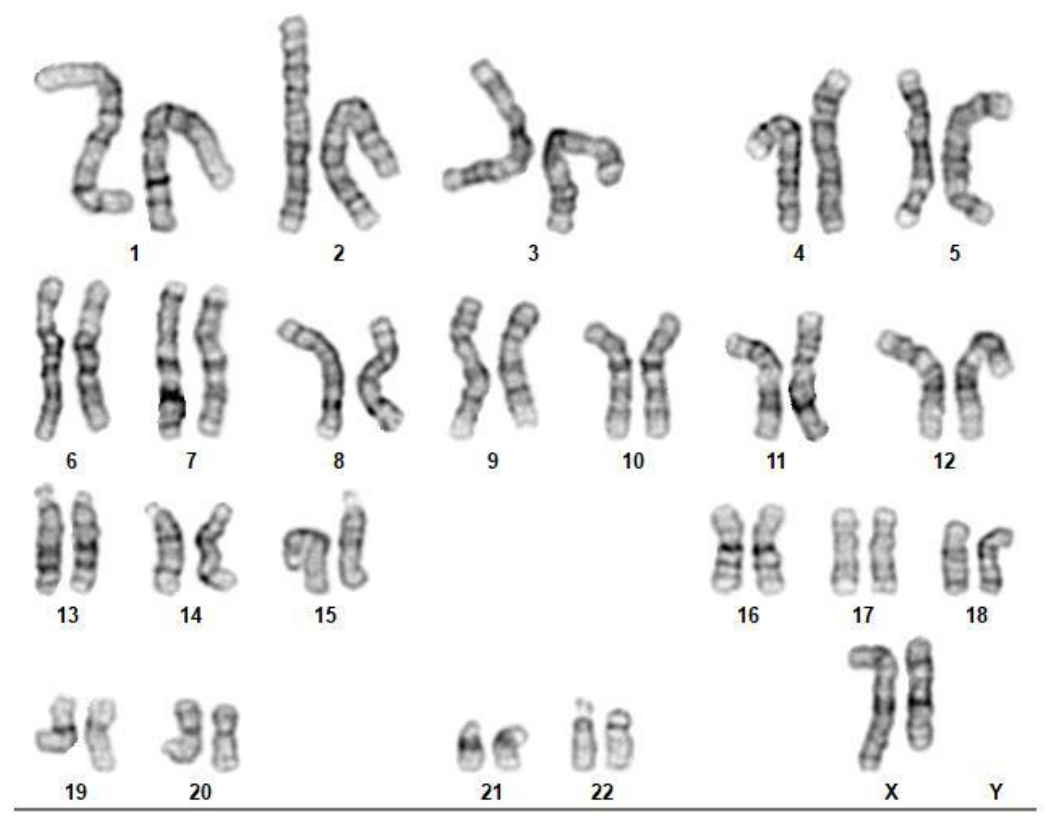

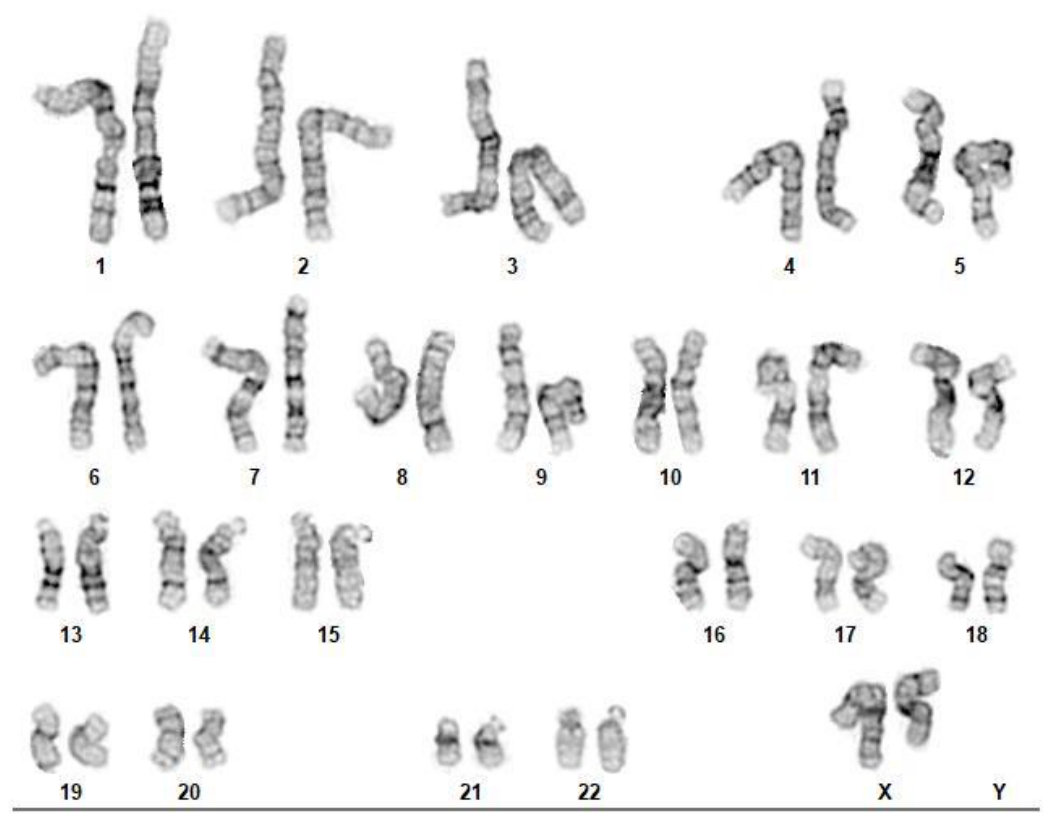

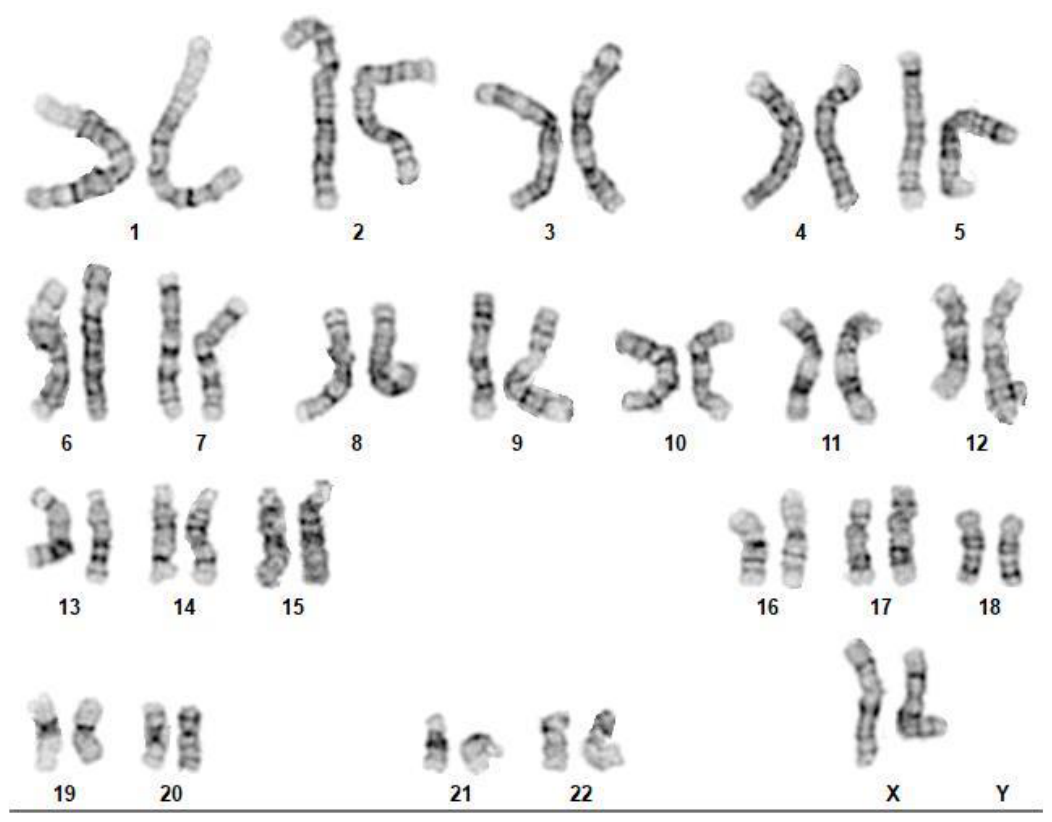

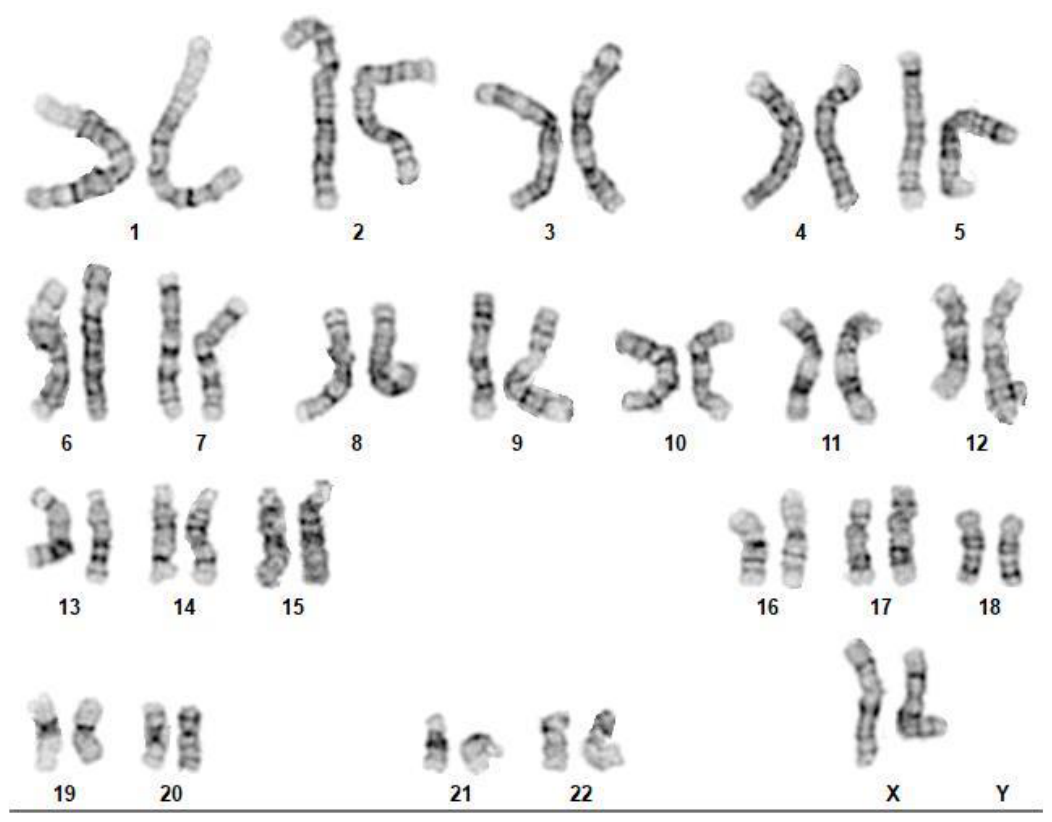

Isogenic

Set A

WT ABCA7

+/- Del ABCA7

-/- Del ABCA7

Isogenic

Set B

Isogenic

Set C

**A**

Supplemental Figure 2.

**Supplemental Figure 2.** Normal karyotypes were observed for all isogenic iPSC lines.
