## Supplemental figure 3 for "Disrupted Lipid Homeostasis as a Pathogenic Mechanism in *ABCA7*-Associated Alzheimer’s Disease Risk"

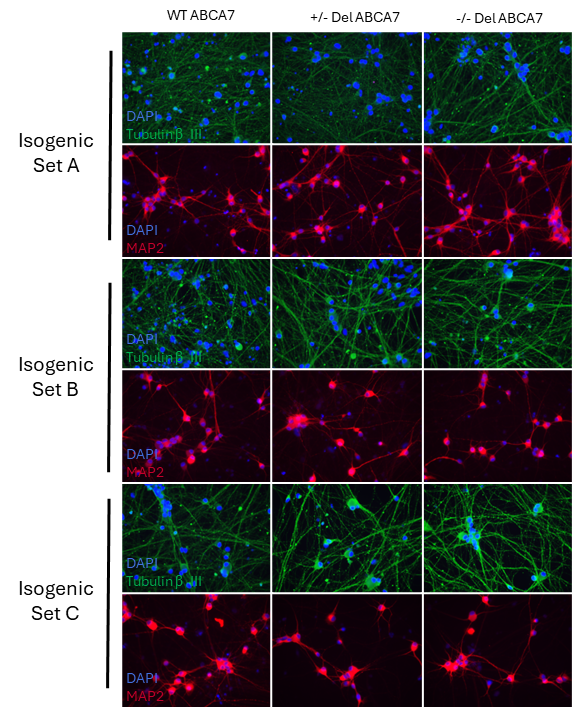

**Supplemental Figure 3.** Neuronal differentiation (iN) using a transdifferentiation approach. iPSC-derived neurons were stained with DAPI, Tubulin beta III, and MAP2 to identify the quality of neuronal differentiation. The transdifferentiation approach produced a homogenous neuronal density in all the isogenic lines.
